# Deep Augmented Multiview Clustering

**DOI:** 10.1101/770016

**Authors:** Ragunathan Mariappan, Jiang Liu, Vaibhav Rajan

**Affiliations:** Department of Information Systems and Analytics, School of Computing, National University of Singapore

## Abstract

We develop a deep learning based method to cluster arbitrary collections of matrices. Our method co-clusters each matrix in the input collection and also associates clusters across matrices thereby enabling discovery of cluster chains. We present preliminary findings on a clinical dataset.

## 1 Introduction

In multiview learning, *views* refer to measurements for the same subjects, that differ in source, datatype or modality. Each matrix, representing a view, has a relationship between two *entity types*, along each matrix dimension, and entity types may be involved in multiple views. See figure 1 for an example. *Collective Matrix Factorization (CMF)* is a general technique to learn shared representations from arbitrary collections of heterogeneous data sources [6]. CMF collectively factorizes the input set of matrices to learn a low-rank latent representation for each entity type from *all* the views in which the entity type is present. It can be used to simultaneously complete one or more matrices in the collection of matrices. Since CMF models arbitrary collections of matrices, this setting is also referred to as *augmented multiview learning* [2].

**Figure 1:**
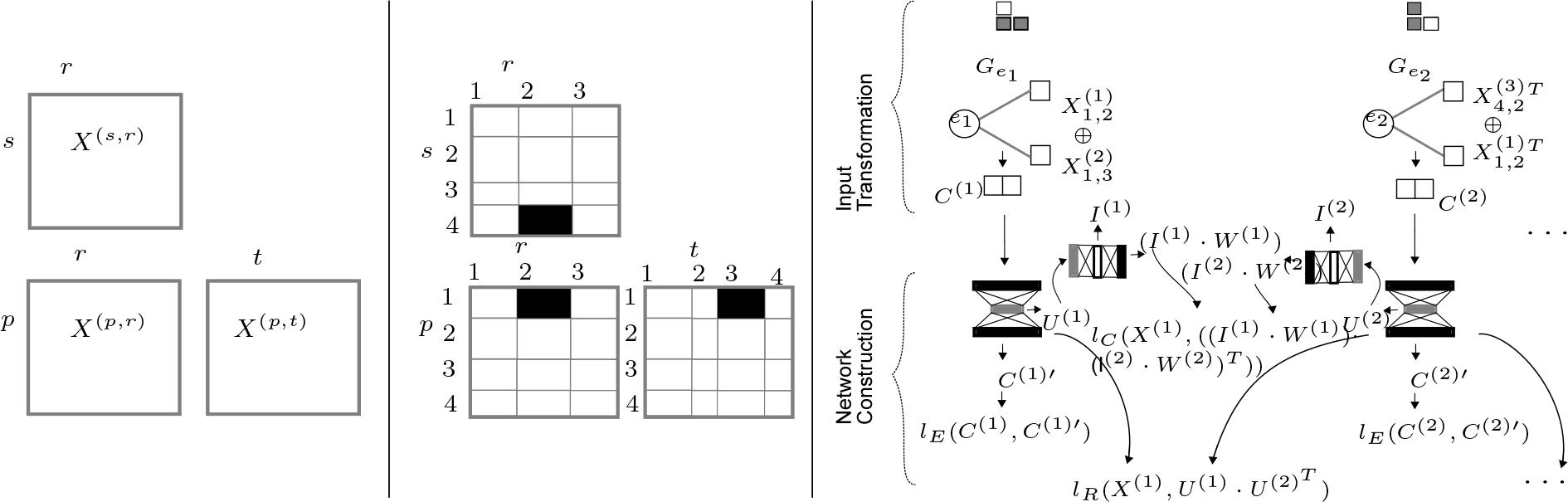
Left: 4 entities *p*: patients, *r*: drugs,*s*: side effect diseases, *t*: treated diseases and 3 relations between the entities, matrices *X*^(*pr*)^, *X*^(*sr*)^, *X*^(*pt*)^, Center: An example entity cluster chain. Here the entities *p*, *r*, *s* and *t* are clustered into 4, 3, 4 and 4 groups respectively. This example captures the cluster chain 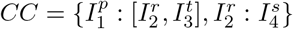, Right: dCMF clustering model construction.

Given *M* matrices (indexed by *m*), 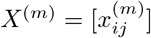, that describe relationships between *E* entities (*e*_1_,…*e*_*E*_), each with dimension *d*_*e*_, our model Deep Collective Matrix Factorization (dCMF) [5] jointly obtains latent representations of each entity 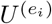 and low-rank factorizations of each matrix 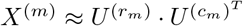, such that 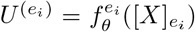 where *f*^(.)^ is an entity-specific non-linear transformation, obtained through a neural network based encoder with weights *θ* and 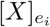 denotes all matrices in the collection that contains a relationship of entity *e*_*i*_. The entities corresponding to the rows and columns of the *m*^*th*^ matrix are denoted by *r*_*m*_ and *c*_*m*_ respectively. There are two steps in dCMF model construction:

1. Input Transformation: For each entity *e*_*i*_, we create a new matrix *C*^(*i*)^, that we call *concatenated matrix*, by concatenating all the matrices containing entity *e*_*i*_.
2. Network Construction: We then use *E* (dependent) autoencoders to obtain the latent factors *U*^(*i*)^ from the concatenated matrices *C*^(*i*)^. For each entity *e*_*i*_ our network has an autoencoder whose input is *C*^(*i*)^, and the decoding is represented by *C*^(*i*)′^. The bottleneck or encoding of each autoencoder, after training, forms the latent factor *U*^(*i*)^. The latent factors are learnt by training all the autoencoders together by solving:

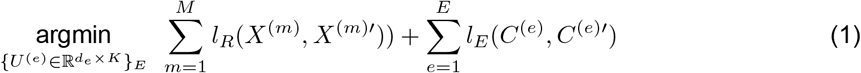

where *l*_*E*_ is the reconstruction loss between the autoencoder’s input *C*^(*i*)^ and the decoding *C*^(*i*)′^; *l*_*R*_ is the matrix reconstruction loss, where the reconstructed matrix 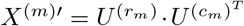 of the view *X*^(*m*)^ is obtained by multiplying the associated row and column entity representations 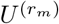 and 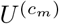.

Collective training of all autoencoders induces dependencies between the autoencoder networks that may result in simultaneous under-fitting in some networks and over-fitting in other networks. We address these optimization challenges through multi-task Bayesian optimization and an acquisition function that is adapted for collective learning of hyperparameters (details in [5]).

### 1.1 Multi-Way Clustering with dCMF representations

In this preliminary study, we extend the dCMF model to obtain, for an arbitrary collection of input matrices, the latent representations of each entity 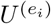 as well as the following:

- disjoint clusters within each entity type, and
- associations between clusters across entity types.

Let 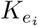 be the number of clusters for entity type *e*_*i*_. To simultaneously learn clusters and associations between them, we assume another factorization for each matrix:

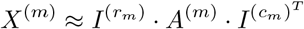

where 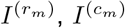 are cluster indicator matrices for the row and column entities respectively, and *A*^(*m*)^ is the cluster association matrix, with *A*_*ij*_ representing the strength of association between cluster *i* of the row entity type and cluster *j* of the column entity type.

The values in 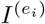 are between 0 and 1, i.e., 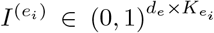 and 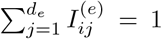, which can be interpreted as a soft cluster assignment score. Further, collective learning of association matrices for all the input matrices enables us to learn multiple *Cluster Chains* that associate clusters across entities in all the input matrices as shown below.

**Illustration.** Figure 1 (left) shows an example of 3 input matrices containing dyadic relationships between 4 entities: patients, drugs, known side effects and treated diseases. Matrices *X*^(*pr*)^*, X*^(*pt*)^ could be extracted from an EMR database and matrix *X*^(*sr*)^ could be from a public repository of drugs and their side effects, e.g., SIDER [3]. Figure 1 (center) shows multi-way clusters in each matrix obtained by clustering each entity. The shaded blocks correspond to one cluster chain found by associating the 4th side effect cluster – the 2nd drug cluster – the 1st patient cluster – the 3rd diseases treated cluster. Similarly other cluster chains exist (not shown) that allow us to associate clusters across entity types for an arbitrary collection of input matrices.

### 1.2 Model Construction

We do not directly tri-factorize the input matrices as done previously in [4, 7]. Instead we use latent representations learnt from dCMF [5] to obtain the cluster indicator and cluster association matrices. This is done by extending the dCMF architecture in two ways:

- **Additional Network Layers.** The autoencoder network for each entity *e*_*i*_ is augmented with a neural network layer for generating 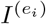 as depicted in Fig 1. A softmax activation ensures that the values in 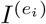 are between 0 and 1, and 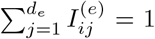. The representation 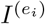 thus generated is passed to another linear layer with no bias to generate *I*^(*e*)^ · *W*^(*e*)^. For a matrix *X*^(*m*)^ with row and column entity representations, 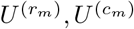, we perform another re-construction through 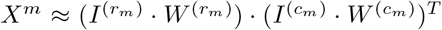. The required tri-factorization can then be obtained by constructing the cluster association matrix as 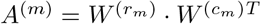.
- **Loss function.** We add another term (*l*_*C*_) to the loss function of dCMF as described below. All the latent factors are learnt by collectively training all the autoencoders by solving:

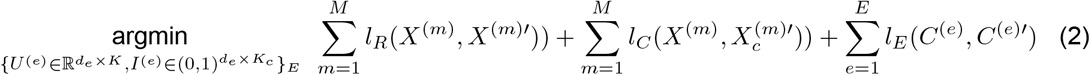

where, in addition to *l*_*E*_ and *l*_*R*_ (from equation 1) we have the matrix reconstruction loss, *l*_*C*_, where the reconstructed matrix 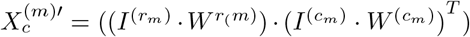 of the view *X*^(*m*)^ is obtained from the associated row and column entity cluster indicators 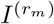 and 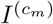.

Note that the cluster association matrix 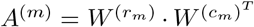 capturing the association between the clusters are obtained by the layer weights. The autoencoders are trained using SGD and Bayesian optimization for hyperparameter tuning as described in [5].

### 1.3 Experimental Results

We assume an augmented multiview setup consisting of three matrices (as shown in Fig 1 (left)).

#### Data

We used the following publicly available data sources:

1. *SIDER*: We construct the matrix *X*^(*s*,*r*)^ using SIDER [3], where the entity types are *s*: known side-effects and *r*: drugs. There are 1322 unique drugs with more than 774K listed side effects.
2. *MIMIC-III*: EMR data of patients were obtained from MIMIC-III [1], and used to construct matrices *X*^(*p*,*r*)^ and *X*^(*p*,*t*)^, with entity types *p*: patients and *t*: diseases-treated. MIMIC-III contains around 651K diagnoses and 4M drug prescriptions data of more than 40K patients.

There were 1322 unique common drugs and 1042 unique common diseases across MIMIC-III and SIDER. Hence we could construct a dataset of the setting in figure 1 with *p* = 39363, *r* = 1322, *s* = 1042, *t* = 1042. The sparsity level of the matrices are *X*^(*p*,*r*)^: 98.8%, *X*^(*s*,*r*)^: 83.3%, *X*^(*p*,*t*)^: 99.5%.

#### Baselines

We compare our results against the baselines Data Fusion by Matrix Factorization (DFMF) [7] and Collective Factorization on Related Matrices (CFRM) [4]. DFMF is a tri-factorization based representation learning method – we use the learnt representations to cluster using K-Means. CFRM is a multi-way spectral clustering algorithm that can be used directly.

#### Evaluation

We create a *test set* of drugs (*r*_*u*_) that exhibit Adverse Drug Reactions (*s*_*u*_) that are hidden during training. They are associated with the corresponding affected patients (*p*_*u*_) and the diseases for which they are treated (*t*_*u*_). We create 4 test sets by randomly selecting 5% of the drugs (twice) to form the test set of drugs and hide, in each case, different percentages (1% and 4%) of their known side effects, We set the number of clusters to be 4 for each of the entities. Starting from the patient clusters, we form chains by finding the associated clusters in the other entities. The methods are evaluated based on the signal found in the cluster chains for these hidden ADRs. This is measured by the *percentage of hidden entities found* in the test set.

#### Results

Tables 1 and 2 show the percentage of hidden entities found in cluster chains from DCMF, DFMF and CFRM. For each cluster chain (in a column) the percentage with respect to each of the entities (along a row) is shown. Results for the other two test sets are similar and not shown.

**Table 1:** 5% drugs randomly selected, 1% of their side-effects hidden; Bold: best results over all clusters in each clustering method, **\***: best result over all clusters in all methods, CC: Cluster Chain.

|  | DCMF |  |  |  | DFMF |  |  |  | CFRM |  |  |  |
| --- | --- | --- | --- | --- | --- | --- | --- | --- | --- | --- | --- | --- |
|  | CC 1 | CC 2 | CC 3 | CC 4 | CC 1 | CC 2 | CC 3 | CC 4 | CC 1 | CC 2 | CC 3 | CC 4 |
| p_%found | 16.66 | 38.71 | 22.42 | 22.22 | 12.74 | 59.98 | 17.42 | 9.86 | 26.29 | 16.39 | 51.66 | 5.66 |
| r_%found | 30.30 | **\*60.61** | 0.00 | 0.00 | 12.12 | **24.24** | 3.03 | **24.24** | **6.06** | 3.03 | **6.06** | 3.03 |
| s_%found | 0.64 | 0.64 | 0.64 | 0.64 | 3.85 | 0.64 | 5.13 | 0.64 | 7.05 | 0.00 | 7.05 | 0.00 |
| t_%found | 3.78 | 3.78 | 3.57 | 10.49 | 2.73 | 7.45 | 67.05 | 15.63 | 7.45 | 3.25 | 1.68 | 7.45 |

**Table 2:** 5% drugs randomly selected, 4% of their side-effects hidden; Bold: best results over all clusters in each clustering method, **\***: best result over all clusters in all methods, CC: Cluster Chain.

|  | DCMF |  |  |  | DFMF |  |  |  | CFRM |  |  |  |
| --- | --- | --- | --- | --- | --- | --- | --- | --- | --- | --- | --- | --- |
|  | CC 1 | CC 2 | CC 3 | CC 4 | CC 1 | CC 2 | CC 3 | CC 4 | CC 1 | CC 2 | CC 3 | CC 4 |
| p_%found | 17.81 | 46.99 | 15.40 | 19.80 | 27.39 | 30.55 | 34.00 | 8.07 | 33.40 | 26.39 | 36.35 | 3.85 |
| r_%found | 3.03 | 9.09 | 3.03 | **\*63.64** | **12.12** | 4.55 | 4.55 | 0.00 | 1.52 | 4.55 | **6.06** | 1.52 |
| s_%found | 1.88 | 3.20 | 1.88 | 1.88 | 3.38 | 6.20 | 3.38 | 3.38 | 8.08 | 8.08 | 8.08 | 1.32 |
| t_%found | 26.25 | 11.46 | 4.58 | 26.25 | 3.02 | 6.88 | 16.46 | 1.46 | 4.38 | 10.10 | 2.08 | 2.50 |

The maximum signal for the hidden entities in our test set is found by one of the chains from DCMF in all our experiments. This suggests that collective learning of representations and cluster structures across the matrices is enabling signal propagation from the diseases found in the EMR to predict side effects that are hidden. Additional experiments are ongoing towards a comprehensive evaluation. In the future, we plan to develop methods to identify the cluster chain of interest by comparing the signal within each chain with known (or predicted) associations from various biomedical ontologies. The representation learning model can be extended by modeling temporal correlations in clinical time series through the use of RNN-based variational autoencoders. Finally, the model can be developed to automatically obtain multiple clusterings in a systematic manner.

## APPENDIX A Deep Learning based Multiview Representations

In this section we provide a short description of Deep Collective Matrix Factorization (dCMF) for Augmented Multiview Learning, a technique developed in the PI’s group at NUS. A complete description can be found in [6]. dCMF is the first deep-learning based method for unsupervised learning of multiple shared representations, that can model non-linear interactions, from an arbitrary collection of matrices.

### A.1 Introduction

Pairwise relational data, found in many domains, can be represented as matrices. Matrix completion, that predicts unknown entries in a matrix, is widely used in many applications, e.g. in recommender systems [5], computer vision [3] and bioinformatics [7], to name a few. Often, the matrices are high-dimensional, sparse, and with inherent redundancies. Sufficient information may be present in latent substructures, that can be approximated through low-rank factorizations, and used in predictive models.

When information from multiple heterogeneous sources is available, predictive models benefit from latent representations that model correlated shared structure. In multi-view learning, *views* refer to measurements for the same subjects, that differ in source, datatype or modality. Each matrix, representing a view, has a relationship between two *entity types*, along each matrix dimension, and entity types may be involved in multiple views. For example, in fig. 2(a), entity *e*_1_ could be patients and clinical data from three different sources (notes *X*^(1)^, images *X*^(2)^, and diagnoses *X*^(3)^) may be used to obtain patient representations for modeling risk of diseases. When auxiliary information about multiple entity types are present, they could be effectively utilized to obtain latent representations. For example, in hybrid recommender systems, where side information matrices about users and movies are used in addition to the historical user-rating matrix to obtain user and movie representations (in fig. 2(b), *X*^(1)^ is the user-rating matrix, *X*^(2)^ has user-features and *X*^(3)^ has movie-features). These latent representations are then used to recommend movies to users.

**Figure 2:**
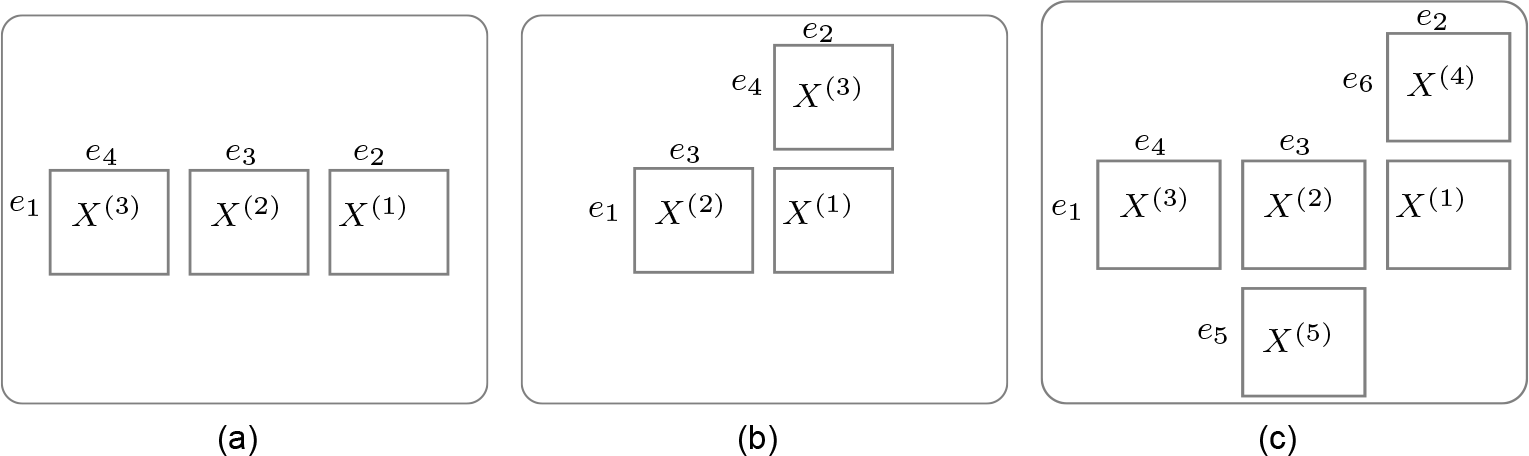
Examples of (a) **multi-view** setting, (b) **recommendation** setting: 4 entities *e*_1_, *e*_2_, *e*_3_, *e*_4_ and 3 relations between the entities, matrices *X*^(1)^, *X*^(2)^, *X*^(3)^; (c) **augmented multi-view** setting: 6 entities *e*_1_, *e*_2_, *e*_3_, *e*_4_, *e*_5_, *e*_6_ and 5 relations between the entities, matrices *X*^(1)^, *X*^(2)^, *X*^(3)^, *X*^(4)^, *X*^(5)^.

*Collective Matrix Factorization (CMF)* is a general technique to learn shared representations from arbitrary collections of heterogeneous data sources [8]. CMF collectively factorizes the input set of matrices to learn a low-rank latent representation for each entity type from *all* the views in which the entity type is present. It can be used to simultaneously complete one or more matrices in the collection of matrices. Since CMF models arbitrary collections of matrices, this setting is also referred to as *augmented multi-view learning* [4]. Fig. 2(c) shows an example. Note that the augmented multi-view setting can generalize to any collection of matrices and subsumes the multi-view and recommendation settings.

Classical matrix factorization based approaches assume linearity in the interaction of latent factors which can be restrictive and fails to capture complex non-linear interactions. Modeling such non-linearities through neural models have significantly improved multi-view learning approaches with two views [11] and multiple (but not augmented) views [10]. A common approach is the use of deep autoencoders to obtain shared representations that form latent factors. However these methods cannot generalize to arbitrary collections of matrices, a limitation that we address in our method.

### A.2 Background

#### Matrix Factorization

For a matrix *X* ∈ ℝ^*m*×*n*^, a low-rank factorization aims to obtain latent factors *U*^(1)^ ∈ ℝ^*m*×*K*^, *U*^(2)^ ∈ ℝ^*n*×*K*^, such that *X* ≈ *U*^(1)^ · *U*^(2)*T*^, where the *K* < min(*m*, *n*) (see fig. 3(a)). The factors are learnt by solving the optimization problem: argmin_*U*,*V*_ *L*(*X*, *U*^(1)^ · *U*^(2)*T*^), where *L* denotes a loss function (e.g., 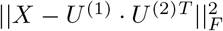, where 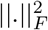 is the Frobenius norm). Collaborative Filtering, for recommendations, uses such an approach where *X* is the rating matrix. A common approach to solving this is through a convex relaxation that minimizes the nuclear norm (the sum of singular values) of 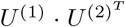, which is equivalent to solving: 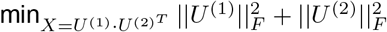 [9].

**Figure 3:**
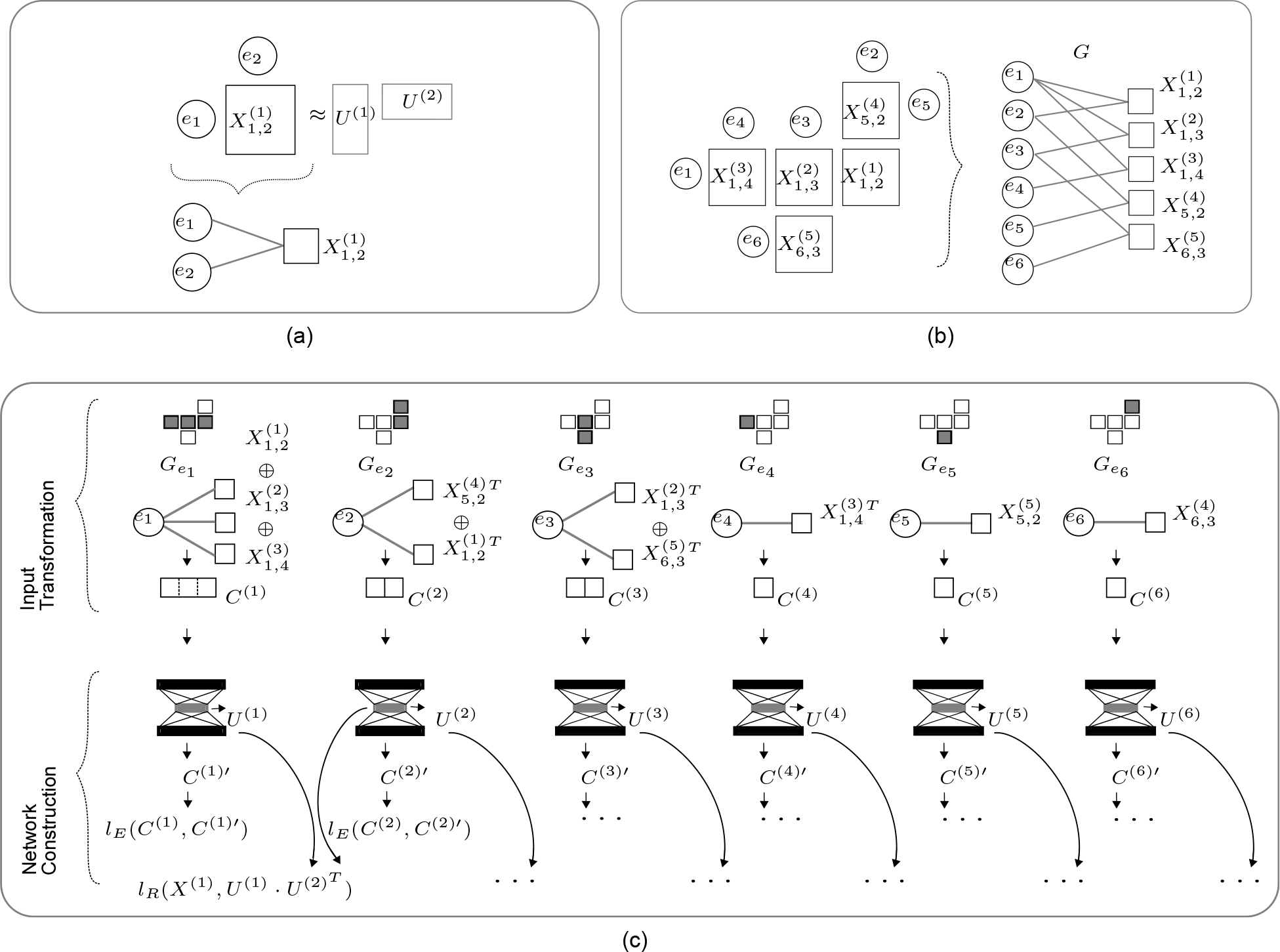
(a) Entity-matrix relationship graph for a single view (b) A collection of views and its entity-matrix relationship graph [square nodes: matrices, circular nodes: entities] (c) dCMF model construction for the example in fig. (b).

**CMF** aims to jointly obtain low-rank factorizations of *M* matrices (indexed by *m*), 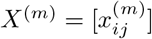, that describe relationships between *E* entities (*e*_1_,…*e*_*E*_), each with dimension *d*_*e*_. The entities corresponding to the rows and columns of the *m*^*th*^ matrix are denoted by *r*_*m*_ and *c*_*m*_ respectively. Fig. 2 shows three examples. Each matrix is approximated by product of low rank-*K* factors that form the representations of the associated row and column entities: 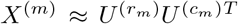 where 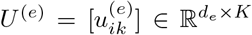 is the low-rank matrix for entity type *e*. Any two matrices sharing the same entity type use the same low-rank representations as part of the approximation, which enables sharing information. For example, in fig. 3(b), the same latent factor 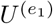 is used to reconstruct the three matrices 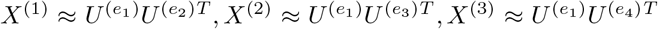. The latent factors are learnt by solving the optimization problem:

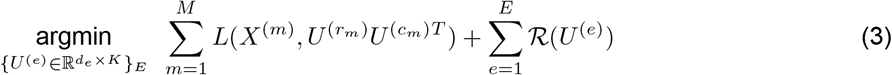

where *M* is the total number of input matrices, *E* is the total number of entities and 𝓡 is a regularizer. For 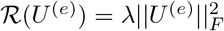, [1] show that this formulation generalizes the nuclear norm for a single matrix to a *collective nuclear norm* defined on an arbitrary set of matrices (with the reasonable assumption that a pair of entity types do not share more than one view). Although this is a non-convex problem, in practice, solutions obtained through Stochastic Gradient Descent yield good performance [1].

### A.3 Deep Collective Matrix Factorization (dCMF)

We design a neural architecture for collective matrix factorization where shared latent representations of each entity are obtained through deep autoencoders. The representations are learnt to enable accurate reconstruction of the input matrices, with the autoencoder reconstruction loss as a regularizer in Eq. 3. Such autoencoder-based regularization has been used in DCCAE [11] for multi-view learning from two views. Our model differs in the way we use autoencoders for learning shared representations. Further, generalizing to augmented multi-view learning leads to non-trivial optimization challenges that we discuss and address in the following sections.

#### Model construction and training

The inputs to dCMF are (1) *M* matrices and (2) the relationship between these matrices and the constituent *E* entities, that can be provided as a bipartite entity-matrix relationship graph *G*(*V*_*E*_, *V*_*M*_, *D*), where vertices *V*_*E*_, *V*_*M*_ represent entities and matrices respectively. Edges (*e*_*i*_, *X*^(*m*)^), (*e*_*j*_, *X*^(*m*)^) ∈ *D* are present if there exists an input matrix *X*^(*m*)^ ∈ *V*_*M*_ capturing the relationship between the entities *e*_*i*_, *e*_*j*_ ∈ *V*_*E*_ (see fig. 3(a),(b)). There are two steps in dCMF model construction:

1. Input Transformation: For each entity *e*_*i*_, we create a new matrix *C*^(*i*)^, that we call *concatenated matrix*, by concatenating all the matrices containing entity *e*_*i*_, i.e., all the matrices that are neighbors of *e*_*i*_ in *G*(*V*_*E*_, *V*_*M*_, *D*). Note that we transform *M* input matrices to *E* concatenated matrices, and a single input matrix (*X*^(*m*)^) may be used in multiple concatenated matrices (*C*^(*i*)^). The concatenation ensures that for each entity, we use the information from all available input matrices to learn its representation.
2. Network Construction: We then use *E* (dependent) autoencoders to obtain the latent factors *U*^(*i*)^ from the concatenated matrices *C*^(*i*)^. For each entity *e*_*i*_ our network has an autoencoder whose input is *C*^(*i*)^, and the decoding is represented by *C*^(*i*)′^. The bottleneck or encoding of each autoencoder, after training, forms the latent factor *U*^(*i*)^. The latent factors are learnt by training all the autoencoders together by solving:

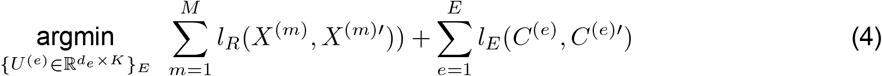

where *l*_*E*_ is the reconstruction loss between the autoencoder’s input *C*^(*i*)^ and the decoding *C*^(*i*)′^; *l*_*R*_ is the matrix reconstruction loss, where the reconstructed matrix 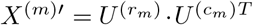 of the view *X*^(*m*)^ is obtained by multiplying the associated row and column entity representations 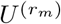 and 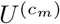. We call the summations in equation (4) the matrix reconstruction loss 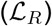 and autoencoder reconstruction loss 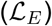 respectively.

Thus, while CMF factorizes each matrix as 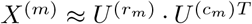, dCMF performs non-linear factorization using autoencoders as 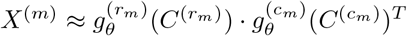, where *g*_*θ*_ is the encoder corresponding to the entity, with parameter set *θ*, obtained by collectively minimizing the sum of all the matrix reconstruction and autoencoder reconstruction losses as described above.

**Illustration.** In fig. 3(a) we show a single matrix 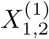 and its two entities *e*_1_ and *e*_2_. The corresponding entity-matrix graph below has 2 circular nodes for two entities and 1 square node for the matrix. In fig. 3(b), we show the graph for the collection of 5 matrices and 6 entities (*e*_1_ to *e*_6_) (from fig. 2(c)). Consider, for instance, the entity *e*_1_. There exists 3 matrices with relationships of entity *e*_1_ with three other entities *e*_2_, *e*_3_ & *e*_4_. Hence there are 3 edges from the node representing *e*_1_ ∈ *V*_*E*_ to the nodes *X*^(1)^*, X*^(2)^*, X*^(3)^ *V*_*M*_.

We illustrate dCMF model construction in fig. 3(c) for the example from fig. 3(b). We construct *E* = 6 autoencoders, one per entity. The autoencoder construction for entity *e*_1_ is illustrated in the first column of fig. 3(c). We show the subgraph 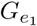 consisting of 3 edges corresponding to the 3 views 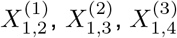. Hence 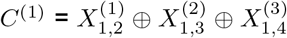, where ⊕ denotes row or columnwise concatenation of the matrices. To pictorially illustrate this we show a miniature of the setup in fig. 3(b) on top of each column in fig. 3(c), greying out the boxes corresponding to the matrices involved in *C*^(*i*)^ construction. We also show *C*^(*i*)^ as a block containing concatenated boxes (equal to the number of matrices *C*^(*i*)^ is composed of) with a label *C*^(*i*)^ below each of the subgraphs which is also the input to the corresponding autoencoder. Similarly we construct the autoencoder for *e*_2_ and the input *C*^(2)^ by concatenating matrices corresponding to the edges of 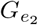 as illustrated in the second column of fig. 3(c). Thus we have 6 columns in fig. 3(c) for each of the 6 entities in setup of fig. 3(b). To avoid clutter in fig. 3(c), we show only two examples of the autoencoder reconstruction loss *l*_*E*_ for entities *e*_1_ and *e*_2_ and one example matrix reconstruction loss *l*_*R*_ for the matrix *X*^(1)^. In total there are *E* = 6 autoencoder reconstruction loss terms and *M* = 5 matrix reconstruction loss terms. Note that this construction can be generalized to any number of entities and matrices.

There are several autoencoder architectures [2] that can be used; we leave that investigation for future work. Here we choose the simplest architecture with multiple fully-connected hidden layers. Notice that the input dimension, which depends on *C*^(*e*)^, is different for each autoencoder and the bottleneck layer dimension (the chosen low rank *K*) is common across all autoencoders. So, the number of layers for each autoencoder is a hyperparameter that is chosen adaptively for each autoencoder.

